# Tooth chipping patterns in *Paranthropus* do not support regular hard food mastication

**DOI:** 10.1101/2021.02.12.431024

**Authors:** Ian Towle, Joel D. Irish, Carolina Loch

## Abstract

The paranthropines, including *Paranthropus boisei* and *Paranthropus robustus*, have often been considered hard-food specialists. The large post-canine teeth, thick enamel, and robust craniofacial features are often suggested to have evolved to cope with habitual mastication of hard foods. Yet, direct evidence for Paranthropus feeding behaviour often challenges these morphological interpretations. The main exception being antemortem tooth chipping which is still regularly used as evidence of habitual mastication of hard foods in this genus. In this study, data were compiled from the literature for six hominin species (including *P. boisei* and *P. robustus*) and 17 extant primate species, to analyse Paranthropus chipping patterns in a broad comparative framework. Severity of fractures, position on the dentition, and overall prevalence were compared among species. The results indicate that both Paranthropus species had a lower prevalence of tooth fractures compared to other fossil hominin species (*P. boisei*: 4%; *P. robustus*: 11%; *Homo naledi*: 37%; *Australopithecus africanus*: 17%; *Homo neanderthalensis*: 45%; Epipalaeolithic *Homo sapiens*: 29%); instead, their frequencies are similar to apes that masticate hard items in a non-regular frequency, including chimpanzees, gibbons, and gorillas (4%, 7% and 9% respectively). The prevalence is several times lower than in extant primates known to habitually consume hard items, such as sakis, mandrills, and sooty mangabeys (ranging from 28% to 48%). Comparative chipping analysis suggests that both Paranthropus species were unlikely habitual hard object eaters, at least compared to living durophage analogues.

## 1. Introduction

When a tooth contacts a hard object with enough force, the enamel can break and fracture to create a chip (Chai and Lawn, 2007; Constantino et al., 2010). The fractured area can vary depending on mechanical and structural properties of the enamel and object (Thomas, 2000; Constantino et al., 2010; He and Swain, 2008; Lawn, Lee, Constantino, and Lucas, 2009; Scott and Winn, 2011). Chips are not generated through a gradual process, although cumulative effects related to enamel fatigue and demineralization may contribute to fracture likelihood and location (Gao et al., 2016; Sanchez-Gonzalez et al., 2020). Importantly, it can take many years of subsequent wear to erase evidence of a chip, making them a crucial marker of past behavior and diet (Belcastro et al., 2018; Towle et al., 2017; Constantino, Markham, and Lucas, 2012).

Enamel chips can be caused by a variety of factors, including food processing, environmental contaminants (e.g., sand or grit), dietary items and social behavior (Constantino et al., 2012; Sauther, Sussman, and Cuozzo, 2002; Scott and Winn, 2011; Stojanowski, Johnson, Paul, and Carver2015; Van Valkenburgh, 2009). These different influences can generate unique chipping patterns across the dentition, and allow inferences of dietary and behavioral factors in archaeological and paleoanthropological samples (Belcastro et al., 2007; Constantino et al., 2010; Scott and Winn, 2011; Nystrom, Phillips-Conroy, and Jolly, 2004; Towle et al., 2017). A range of recent human populations, fossil hominins and extant primates have been studied for evidence of chipping (e.g., Belcastro et al., 2007; Bonfiglioli et al., 2004; Gould, 1968; Lous, 1970; Molnar et al., 1972; Silva, Gil, Soares, and da Silva, 2016; Turner and Cadien, 1969; Constantino et al., 2010; Grine et al., 2010; Robinson, 1954; Tobias, 1967; Johanson and Taieb, 1976; Ward et al., 2001; Fox and Frayer, 1997; Scott and Winn, 2011; Stojanowski et al., 2015; Larsen, 2015; Lous, 1970; Molnar et al., 1972; Fannin et al., 2020). However, until recently, differences in recording methods have made inter-study comparisons challenging.

Enamel chipping may provide critical insights into the feeding habits of enigmatic hominins, such as the ‘robust Australopith’ clade, e.g., *Paranthropus boisei* and *P. robustus. Paranthropus* species have long been inferred to be hard object feeders (durophagous), with large bunodont posterior teeth and robust cranial features thought to reflect adaptations related to habitually masticating hard foods (Rak, 1983; Jolly, 1970; Teaford and Ungar, 2000; Constantino et al., 2018). However, the diets of these species are still debated, with a variety of dietary scenarios described (e.g., Martínez et al., 2016; Williams, 2015; Strait et al., 2013; Smith et al., 2015; Cerling et al., 2011; Van der Merwe et al., 2008; Wood and Strait, 2004). Additionally, there is often an apparent disconnect between diet and craniodental morphology in *Paranthropus*.

Direct evidence from enamel microwear studies of *P. boisei* suggests little to no hard object feeding (Ungar et al., 2008). In *P. robustus*, higher enamel surface complexity may indicate hard foods were consumed more frequently (Ungar, 2019; Scott et al., 2005; Peterson et al., 2018), potentially in the context of ‘fallback foods’. However, the role of hard plant tissues in generating microwear features is currently debated (van Casteren et al., 2020; Teaford et al., 2020). From stable carbon isotopes, the diets of *P. robustus* and *P. boisei* appear substantially different (Cerling et al., 2011; Ungar and Sponheimer, 2011). Isotopic results for *P. boisei* are in concordance with microwear and some biomechanical evidence, suggesting that hard foods such as seeds and nuts did not make up a significant part of their diet; rather, C4 graminoids (e.g., grasses and sedges) were likely common food sources (Guatelli-Steinberg, 2016; Macho, 2014; Dominy et al., 2008; Yeakel et al., 2007; Cerling et al., 2011; Kaiser et al., 2019). A more recent study on comparative biomechanical and morphological data in primates also suggests a soft-food niche for *Paranthropus* (Marcé-Nogué et al., 2020). Based on this and other evidence, many dental anthropologists now regard *Paranthropus* as non-hard food eaters (Ungar and Hlusko, 2016; Grine and Daegling, 2017; Kaiser et al., 2019), while others consider them dietary generalists (e.g., eurytopy; Wood and Strait, 2004; Strait et al., 2013).

Nonetheless, some researchers maintain that hard foods were commonly consumed in *Paranthropus*, and/or played a significant role in the craniodental evolution of the genus (e.g., Constantino et al., 2009; Constantino et al., 2018; Paine et al., 2019; Strait et al., 2013; Smith et al., 2015). Chipping is usually one of the main forms of evidence proposed to support these hypotheses, typically to show that hard object consumption was common (Constantino et al., 2010, Constantino et al., 2018; Ungar, 2019; Paine et al., 2019). Earlier work suggested that ingestion of grit or bones may instead be responsible for the chipping in their teeth (Robinson, 1954; Tobias, 1967).

Despite being crucial in elucidating the evolution and diet of *Paranthropus*, a broad comparison specifically focusing on chipping patterns in *Paranthropus* relative to other hominins and extant primates is still lacking. In particular, although data in the current article comes from published sources, because *Paranthropus* was not the focus of these studies, it has led to misunderstandings of interpretation. For example, Paine et al. (2019:104) stated that *P. robustus* had a “significant degree of hard object feeding,” with Towle et al. (2017) cited to support this claim. In reality, Towle et al. (2017) reported that *P. robustus* has the lowest rate of chipping of any hominin studied, with a prevalence similar to gorillas. Additionally, in previous studies *Paranthropus* species are often only compared to one another, or to other fossil hominins. For example, the *P. robustus* diet is said to have contained a “significant amount of hard food content,” based on higher chipping prevalence than in *P. boisei* (Constantino et al., 2018:76). Therefore, chipping is often considered evidence of regular hard food consumption, but extant comparisons are needed to determine if these conclusions are supported in primates that are known hard-object feeders.

In three recent studies that include comparisons of chipping in a total of 25 extant primate species (Towle and Loch, 2021; Fannin et al., 2020; Towle et al., 2017), all samples show at least some chips. Therefore, the presence of chipping in a sample on its own may tell us little about diet or behavior. Instead, the evaluation of the patterns in extant primates with associated ecological data allows an understanding of chipping relative to diet and behavior. In this study, we compared these patterns in a range of extant primates and fossil hominins, to test whether prevalence and patterns in *Paranthropus* support habitual, or occasional, durophagy. The extant primates analyzed include several species considered hard-object feeding specialists (sooty mangabeys, mandrills and sakis). Other species either focus on particular non-hard food items or have been reported to rarely consume hard foods (including several ape and colobus species), and those with a more varied diet, including eurytopy and terrestrial species (e.g., Japanese macaques, mandrills and baboons). As well as prevalence, the size and distribution of chips across the dentition when compared to other species may help elucidate behavioral factors that led to their occurrence in *Paranthropus*.

## 2. Materials and Methods

Data were compiled from the recent literature. Fossil hominin samples include specimens assigned to *Homo naledi, Australopithecus africanus, P. robustus, P. boisei*, and *H., neanderthalensis* (following Towle et al., 2017; Constantino et al., 2018; Constantino and Lawn, 2019; Belcastro et al., 2018). Extant primates include chimpanzees (*Pan troglodytes troglodytes*), Western lowland gorillas (*Gorilla gorilla gorilla*), Kloss’s gibbons (*Hylobates klossii*), hamadryas baboons (*Papio hamadryas*), pig-tailed langurs (*Simias concolor*),Japanese macaques (*Macaca fuscata*), Dent’s mona monkeys (*Cercopithecus denti*), blue monkeys (*Cercopithecus mitis*), mandrills (*Mandrillus leucophaeus* and *M. sphinx*), Raffles’ banded langurs (*Presbytis femoralis*), Mentawai langurs (*P. potenziani*), and brown woolly monkeys (*Lagothrix lagothricha*), Sakis (*Pithecia* sp.), sooty mangabeys (*Cercocebus atys*),olive colobus monkeys (*Procolobus verus*), Western red colobus (*Piliocolobus badius*) and king colobus (*Colobus polykomos;* following Towle and Loch, 2021; Towle et al., 2017; Fannin et al., 2020). In these studies, data collection methods were standardized, with only minor variations in techniques. An exception is Fannin et al. (2020), where only fourth premolars and first molars were studied. A summary of methods is given below.

Some hominin species, particularly *P. robustus and A. africanus*, were studied previously for chipping prevalence (e.g., Grine et al., 2010; Robinson, 1956; Wallace, 1973; Tobias, 1967). Here, data from Towle et al. (2017) are used for these species. Teeth with minimal wear were removed for prevalence in Towle et al., (2017), but are included here to allow direct comparisons with other samples, resulting in slightly different frequencies than before.

Antemortem fractures were only recorded if subsequent attrition is evident on the chipped surface (i.e., the chip scar), to rule out postmortem damage (Scott and Winn, 2011; Belcastro et al., 2018). Smoothing and coloration were used for this purpose, i.e., postmortem fractures displaying ‘fresh’ enamel brighter than the rest of the crown and with sharp edges (Towle and Loch, 2021). The total number of chips on each tooth was also recorded for some samples (Belcastro et al., 2018; Towle et al., 2017; Towle and Loch, 2021). Results refer to all permanent teeth unless stated otherwise.

Fractures were recorded on a three-point grading system following either Bonfiglioli et al. (2004) or Towle and Loch (2021). The latter uses slight modifications that remove direct measurements to allow a broader range of comparisons among primates:

1. Small enamel chip (crescent-shaped) on the outer edge of the enamel. Dentine is not exposed and the chip is restricted to the outer rim of the occlusal surface.
2. Larger chip that extends near to the enamel-dentine junction. A small area of dentine might be exposed.
3. Large irregular fracture in which a significant area of dentine is exposed. More dentine than enamel was removed by the fracture.

## 3. Results

Prevalence and overall patterns of permanent tooth chipping for each species with comparable data are summarized in Table 1. Hominins cover the range of prevalence in extant primates, with chipping common in *Homo* (37–45% of all permanent teeth) and rare in *Paranthropus* (4–11%). *Australopithecus africanus* falls between these two extremes with 17% of teeth displaying at least one fracture. The prevalence in *Paranthropus* is similar to the extant ape species studied, chimpanzees, gibbons and gorillas (Table 1). The chipping rate in extant primates considered to be hard-food specialists (e.g., sakis and mandrills, 28% and 37% respectively) is several times greater than in both *Paranthropus* species (*P. boisei*,4.4%; *P. robustus*, 11.1%; Table 1).

**Table 1.** Prevalence of chipping in different primate species, by tooth type, jaw, and size of chips. Data refers to all permanent teeth unless stated.

| Species | Common name | Posterior teeth | Anterior teeth | All teeth % | Multiple chipped teeth % | Small:large chip ratio | Reference |
| --- | --- | --- | --- | --- | --- | --- | --- |
| *Homo naledi* |  | 46.39 (45/97) | 20.75 (11/53) | 37.33 (56/150) | 50.00 | 8.33 | Towle et al. (2017) |
| *Australopithecus africanus* |  | 20.16 (50/248) | 7.32 (6/82) | 16.97 (56/330) | 16.07 | 10.20 | Towle et al. (2017) |
| *Paranthropus robustus* |  | 10.90 (23/211) | 11.86 (7/59) | 11.11 (30/270) | 6.67 | 1.73 | Towle et al. (2017) |
| *Paranthropus boisei*<sup>a</sup> |  | — | — | 4.4<sup>b</sup> | — | — | Constantino and Lawn (2019) |
| *Homo neanderthalensis* | Neanderthals | 31.53 (35/111) | 64.86 (48/74) | 44.86 (83/185) | 9.00 | 0.90 | Belcastro et al. (2018) |
| *Gorilla gorilla gorilla* | Western lowland gorilla | 11.70 (137/1171) | 5.25 (32/610) | 9.49 (169/1781) | 4.14 | 10.27 | Towle et al. (2017) |
| *Pan troglodytes* | Chimpanzee | 4.52 (65/1439) | 4.13 (33/800) | 4.38(98/2239) | 2.04 | 2.27 | Towle et al. (2017) |
| *Cercopithecus denti* | Dent's mona monkey | 16.11 (29/180) | 27.06 (23/85) | 19.62 (52/265) | 1.92 | 5.50 | Towle and Loch (2021) |
| *Cercopithecus mitis* | Blue monkey | 18.13 (29/160) | 9.76 (8/82) | 15.29 (37/242) | 13.51 | 6.40 | Towle and Loch (2021) |
| *Mandrillus spp.* | Mandrill | 36.11 (39/108) | 40.00 (8/20) | 36.72 (47/128) | 23.40 | 6.83 | Towle and Loch (2021) |
| *Presbytis femoralis* | Raffles' banded langur | 31.88 (51/160) | 40.24 (33/82) | 34.71 (84/242) | 11.90 | 4.60 | Towle and Loch (2021) |
| *Presbytis potenziani* | Mentawai langur | 20.25 (32/158) | 17.39 (12/69) | 19.38 (44/227) | 11.36 | 43.00 | Towle and Loch (2021) |
| *Hylobates klossii* | Kloss's gibbon | 5.60 (13/232) | 11.90 (10/84) | 7.28 (23/316) | 0.00 | 1.88 | Towle and Loch (2021) |
| *Macaca fuscata* | Japanese macaque | 24.68 (171/693) | 15.33 (44/287) | 21.94 (215/980) | 12.56 | 5.76 | Towle and Loch (2021) |
| *Simias concolor* | Pig-tailed langur | 16.61 (46/277) | 19.70 (26/132) | 17.60 (72/409) | 12.50 | 7.00 | Towle and Loch (2021) |
| *Lagothrix lagothricha* | Brown woolly monkey | 21.79 (34/156) | 11.27 (8/71) | 18.50 (42/227) | 0.00 | 7.40 | Towle and Loch (2021) |
| *Pithecia spp.* | Saki | 35.09 (60/171) | 11.59 (8/69) | 28.33 (68/240) | 23.53 | 5.80 | Towle and Loch (2021) |
| *Papio hamadryas* | Hamadryas baboon | 26.49 (89/336) | 9.89 (18/182) | 20.66 (107/518) | 8.41 | 7.23 | Towle and Loch (2021) |

<sup>a</sup>Includes deciduous teeth.
<sup>b</sup>Sample size not provided.

Further direct comparisons are made with additional species from Fannin et al. (2020) for first molars and fourth premolars in Figure 1, with *P. robustus* displaying the fourth lowest rate of fractures out of 20 species studied. Further, this equates to a chipping prevalence for these teeth approximately five times fewer than both Sooty mangabey and *H. naledi* (Figure 1). When split into individual tooth types, all *P. robustus* teeth show consistently low chipping rates, except for canines and third premolars, with a moderate prevalence (21% and 23%respectively). Each *P. robustus* molar type (first, second and third) shows a low prevalence of chipping, with each displaying one of the lowest rates relative to the same tooth in other species (Table 2). Few *P. robustus* teeth exhibit multiple chips (6.7%of chipped teeth have more than one fracture). *Paranthropus robustus* also has a high ratio of larger chips relative to most other species (1.7 small chip for every large fracture); only chimpanzees, gibbons and Neanderthals have similar proportions of large chips (2.3, 1.9 and 0.9 small for every large fracture, respectively). When split into tooth categories, large chips were more frequent on anterior teeth and premolars in *P. robustus*, with molars showing a low rate of larger fractures similar to other species (Table 3).

**Figure 1.**
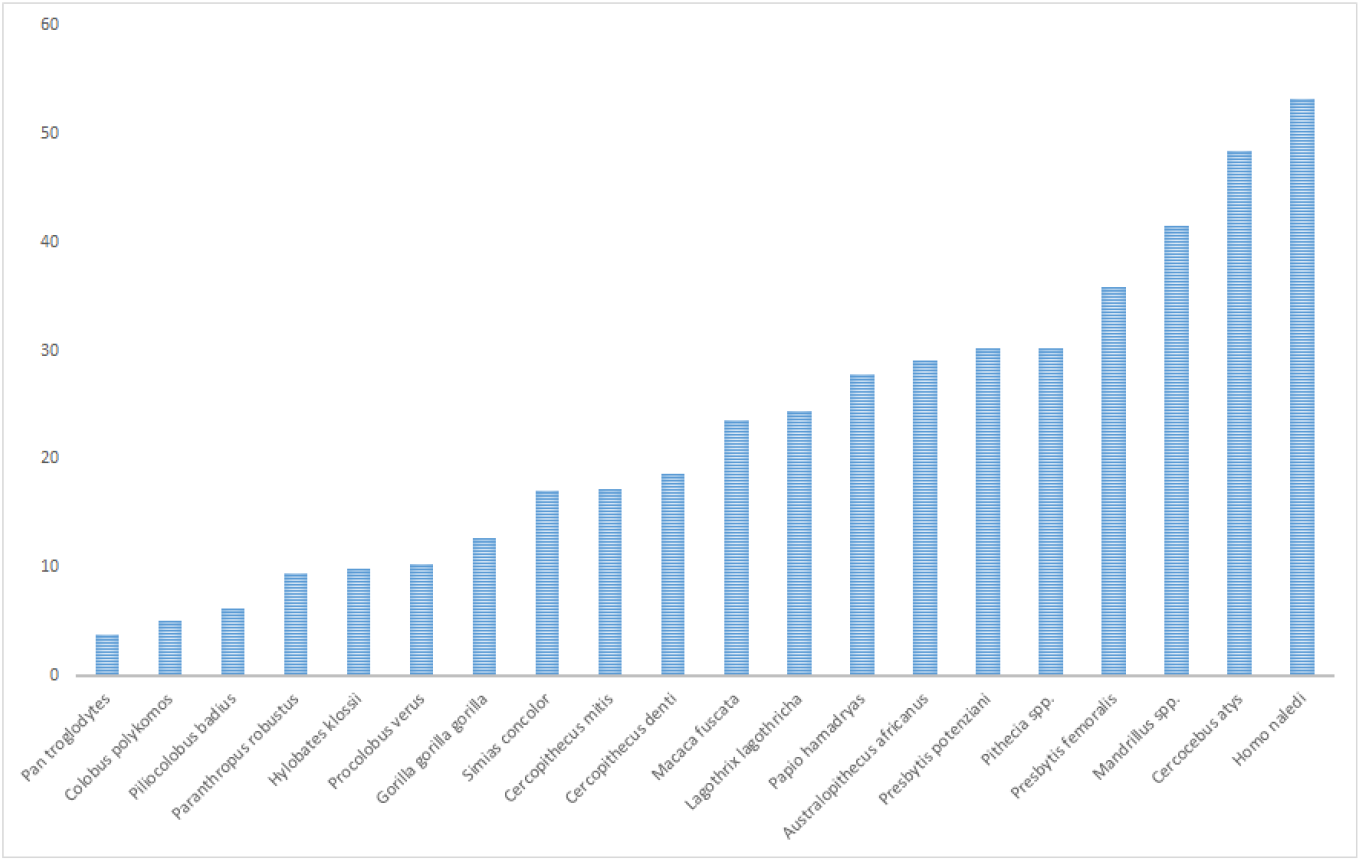
Chipping prevalence for first molars and fourth premolars for different extant primate and fossil hominin species. Species are organized by increasing chipping prevalence.

**Table 2.** Prevalence of chipping (%) in different primate species, split by individual permanent tooth types. I: incisor; C: canine; PM: premolar; M: molar.

| Species | Common name | I1 | I2 | C | PM3 | PM4 | M1 | M2 | M3 |
| --- | --- | --- | --- | --- | --- | --- | --- | --- | --- |
| *Homo naledi* |  | 40 (6/15) | 12.5 (2/16) | 13.64 (3/22) | 42.11 (8/19) | 42.11 (8/19) | 60.71 (17/28) | 36.84 (7/19) | 41.67 (5/12) |
| *Australopithecus africanus* |  | 17.39 (4/23) | 4.35 (1/23) | 2.78 (1/36) | 18.60 (8/43) | 32.43 (12/37) | 29.53 (13/49) | 17.91 (12/67) | 9.62 (5/52) |
| *Paranthropus robustus* |  | 9.38 (3/32) | 7.69 (1/13) | 21.43 (3/14) | 23.33 (7/30) | 4.65 (2/43) | 13.46 (7/52) | 4.65 (2/43) | 11.63 (5/43) |
| *Gorilla gorilla gorilla* | Western lowland gorilla | 4.05 (9/222) | 5.12 (11/215) | 6.94 (12/173) | 3.76 (8/213) | 9.26 (20/216) | 15.22 (44/289) | 10.89 (28/257) | 18.88 (37/196) |
| *Pan troglodytes* | Chimpanzee | 2.79 (8/287) | 4.06 (11/271) | 5.79 (14/242) | 2.22 (6/270) | 1.87 (5/268) | 5.18 (20/386) | 3.46 (10/289) | 10.62 (24/226) |
| *Cercopithecus denti* | Dent's mona monkey | 38.71 (12/31) | 34.62 (9/26) | 7.14 (2/28) | 8.57 (3/35) | 10.81 (4/37) | 26.32 (10/38) | 15.79 (6/38) | 18.75 (6/32) |
| *Cercopithecus mitis* | Blue monkey | 17.86 (5/28) | 4 (1/25) | 6.90 (2/29) | 3.13 (1/32) | 15.63 (5/32) | 18.75 (6/32) | 25 (8/32) | 28.13 (9/32) |
| *Mandrillus spp.* | Mandrill | 55.56 (5/9) | 50 (3/6) | 0 (0/5) | 13.64 (3/22) | 33.33 (7/21) | 50 (10/20) | 50 (11/22) | 34.78 (8/23) |
| *Presbytis femoralis* | Raffles' banded langur | 52.17 (12/23) | 46.15 (12/26) | 34.48 (10/29) | 18.75 (6/32) | 31.25 (10/32) | 40.63 (13/32) | 31.25 (10/32) | 37.5 (12/32) |
| *Presbytis potenziani* | Mentawai langur | 18.18 (4/22) | 22.22 (6/27) | 10 (2/20) | 6.45 (2/31) | 31.25 (10/32) | 29.03 (9/31) | 31.25 (10/32) | 3.13 (1/32) |
| *Hylobates klossii* | Kloss's gibbon | 2.78 (1/36) | 12.5 (4/32) | 31.25 (5/16) | 4.26 (2/47) | 6.52 (3/46) | 13.33 (6/45) | 3.85 (2/52) | 0 (0/42) |
| *Macaca fuscata* | Japanese macaque | 22.62 (19/84) | 16.28 (14/86) | 9.40 (11/117) | 17.91 (24/134) | 18.66 (25/134) | 27.92 (43/154) | 25.17 (36/143) | 33.59 (43/128) |
| *Simias concolor* | Pig-tailed langur | 29.79 (14/47) | 13.33 (6/45) | 15 (6/40) | 8.62 (5/58) | 10.53 (6/57) | 24.07 (13/54) | 28.07 (16/57) | 11.76 (6/51) |
| *Lagothrix lagothricha* | Brown woolly monkey | 10.53 (2/19) | 11.11 (3/27) | 15.79 (3/19) | 15.38 (4/26) | 25.93 (7/27) | 22.22 (4/18) | 22.22 (6/27) | 33.33 (6/18) |
| *Pithecia spp.* | Saki | 13.33 (2/15) | 16.67 (4/24) | 8.70 (2/23) | 26.67 (8/30) | 24.14 (7/29) | 37.5 (9/24) | 44.44 (12/27) | 50 (13/26) |
| *Papio hamadryas* | Hamadryas baboon | 8.47 (5/59) | 9.68 (6/62) | 11.48 (7/61) | 15.87 (10/63) | 17.65 (12/68) | 37.5 (27/72) | 37.68 (26/69) | 21.88 (14/64) |

**Table 3.** Prevalence of large chips, split by tooth type. Number of large chips (severity 2 and 3) as a percentage of total chips.

| Species | Common name | Molars | Premolars | Canines | Incisors |
| --- | --- | --- | --- | --- | --- |
| *Paranthropus robustus* |  | 14.29 (2/14) | 44.44 (4/9) | 100 (3/3) | 50 (2/4) |
| *Homo naledi* |  | 17.24 (5/29) | 6.25 (1/16) | 0 (0/3) | 0 (0/8) |
| *Australopithecus africanus* |  | 6.67 (2/30) | 10 (2/20) | 100 (1/1) | 0 (0/5) |
| *Gorilla gorilla gorilla* | Western lowland gorilla | 7.34 (8/109) | 10.71 (3/28) | 0 (0/12) | 20 (4/20) |
| *Chimpanzees* | Chimpanzee | 35.19 (19/54) | 27.27 (3/11) | 42.86 (6/14) | 10.53 (2/19) |
| *Cercopithecus denti* | Dent's mona monkey | 22.73 (5/22) | 0 (0/7) | 0 (0/2) | 14.29 (3/21) |
| *Cercopithecus mitis* | Blue monkey | 8.70 (2/23) | 33.33 (2/6) | 0 (0/2) | 16.67 (1/6) |
| *Mandrillus spp.* | Mandrill | 17.24 (5/29) | 10 (1/10) | n/a (0/0) | 0 (0/8) |
| *Presbytis femoralis* | Raffles' banded langur | 8.57 (3/35) | 37.5 (6/16) | 30 (3/10) | 12.5 (3/24) |
| *Presbytis potenziani* | Mentawai langur | 5 (1/20) | 0 (0/12) | 0 (0/2) | 0 (0/10) |
| *Hylobates klossii* | Kloss's gibbon | 50 (4/8) | 20 (1/5) | 40 (2/5) | 20 (1/5) |
| *Macaca fuscata* | Japanese macaque | 22.13 (27/122) | 2.04 (1/49) | 18.18 (2/11) | 3.03 (1/33) |
| *Simias concolor* | Pig-tailed langur | 11.43 (4/35) | 9.09 (1/11) | 0 (0/6) | 20 (4/20) |
| *Lagothrix lagothricha* | Brown woolly monkey | 18.75 (3/16) | 0 (0/11) | 33.33 (1/3) | 0 (0/5) |
| *Pithecia spp.* | Saki | 8.82 (3/34) | 26.67 (4/15) | 0 (0/2) | 0 (0/6) |
| *Papio hamadryas* | Hamadryas baboon | 11.94 (8/67) | 9.09 (2/22) | 14.29 (1/7) | 18.18 (2/11) |

## 4. Discussion and conclusions

Chipping prevalence in extant primates links well with dietary and behavioral observations. Hard object-feeding primates have a high prevalence of tooth chipping (Towle and Loch, 2021; Fannin et al., 2020). Species considered hard-object feeding specialists have a prevalence >25%, with the diets of sooty mangabeys, mandrills and sakis containing significant amounts of hard foods (e.g., durophagy; Kinzey and Norconk, 1993; Fleagle and McGraw, 1999; McGraw et al., 2011; Pampush et al., 2013; Fannin et al., 2020; vanCasteren et al., 2020). The overall chipping prevalence in both *Paranthropus* species is far below this threshold (4-11%). Furthermore, the prevalence in first molars and fourth premolars (for which most data is currently available) shows *P. robustus* with one of the lowest chipping rates.

Based on these results, the current study indicates that both *Paranthropus* species experienced significantly fewer crown chips than other hominin species, and several times less than extant primates consuming hard objects (e.g., food items and/or grit). The chipping prevalence in *Paranthropus* is similar to species that rarely masticate hard items, including several apes, colobines, and guenons. The findings do not corroborate that either *Paranthropus* species were habitual hard-object feeders, at least to the extent of modern durophagous primates such as sakis, mandrills, and sooty mangabeys. Daegling et al., (2013) suggested that the large posterior teeth of *Paranthropus* showed a greater prevalence of chips due to greater surface area. Crown fractures are also more likely to form on teeth with enamel defects or unusual wear (Soukup, 2019). The pitting enamel hypoplasia present on a large proportion of *Paranthropus* molars (Towle and Irish, 2019) suggests they may have been more fracture prone. Thus, durophagy may have been even less frequent than the low chipping rate in both *Paranthropus* species suggests. The low rate of multiple chips on a single tooth supports this conclusion.

There are other factors that influence chipping prevalence. In humans, chips are generally more common on anterior teeth due to food processing, trauma, or non-masticatory cultural behavior, but frequencies vary substantially among groups (Scott and Winn, 2011; Stojanowski et al., 2015; Bonfiglioli et al., 2004; Gould, 1968; Larsen, 2015; Lous, 1970; Molnar et al., 1972). Grit is also masticated by many primates, and likely influences chipping rates in certain species (e.g., Van Casteren et al., 2019; Fannin et al., 2020; Towle et al., 2017). Both of these factors are associated with an increase in chipping frequency. Therefore, a low rate of chipping in *Paranthropus* likely suggests that trauma related to both food consumption and other factors such as grit mastication were rare.

A possible exception is the moderate rate of large fractures on canines and third premolars, which may suggest a modest level of trauma in these regions. The average size of these fractures is similar to those on Neanderthal anterior teeth (Belcastro et al., 2018). However, a low rate of chipping on *P. robustus* incisors suggests a non-masticatory explanation is unlikely. Similarly, grit tends to create smaller fractures, which are more evenly distributed over the posterior dentition (Towle et al., 2017); thus, grit is also an unlikely cause for chipping in *P. robustus*. One possible explanation could be specific masticatory behaviors, such as placing hard foods in these positions for the initial phase of mastication (e.g., breaking seeds or nuts). That said, the small sample size for individual tooth types makes inferences difficult, and the chipping rate is still only moderate when compared to other species. Therefore, although it is possible that *P. robustus* did occasionally masticate hard foods, there is little evidence to suggest this was a common practice.

Constantino et al. (2018) reported a rate of 5% chipping for *P. robustus*, which is similar to a more recent finding of 4.4% of teeth in *P. boisei* (Constantino and Lawn, 2019). The difference in prevalence for *P. robustus* in Towle et al. (2017) likely relates to stricter criteria for inclusion in the former study (i.e., deciduous and postmortem damaged teeth were removed from analysis), with both studies recording approximately 30 chipped teeth. If all teeth are included from the Towle et al (2017) data set (i.e., including postmortem damaged teeth) there is a more similar chipping rate of 7.46%. Therefore, the chipping prevalence in *Paranthropus* seems consistently low across studies when compared to other primates, and the actual chipping prevalence between *Paranthropus* species is likely similar.

It has been suggested that *Paranthropus* preferred soft or tough foods, but relied on harder foods as ‘fallback foods’ (Constantino and Wright, 2009; Ungar et al., 2008). This hypothesis suggests *Paranthropus* evolved large teeth and robust cranial structures to cope with dietary items that were rarely consumed, but required requisite adaptations. This hypothesis has been challenged, with other scenarios favored (Daegling et al., 2013; Scott et al., 2014). Extant primate chipping patterns may offer some insight. Species that evolved specialized dental characteristics for softer foods can still occasionally eat hard foods (e.g., van Casteren et al., 2019). Additionally, an occasional hard object-feeding primate (consumed as seasonal or as fallback foods) can show elevated chipping frequencies. For example, while the diet of brown woolly monkeys is primarily based on soft fruits, at certain times of the year they consume seeds and hard fruits (Peres, 1994; Defler and Defler, 1996); this dietary shift could explain their relatively high chipping prevalence, although further research is needed into the mechanical properties of these foods.

The orientation of the hard object and underlying enamel microstructure is crucial in chip formation, as are the size and shape of the tooth, the object, and resultant biomechanical forces (Xu et al., 1998; Lucas et al., 2008; Chai and Lawn, 2007). As such, species-specific enamel attributes (e.g., thickness and mechanical properties) likely evolved for functional reasons (Cuy et al., 2002); species that regularly eat hard foods likely evolved dental characteristics in response to high biomechanical demands (Ungar and Lucas, 2010). It has been suggested that thick enamel may have evolved in some hominins, including *Paranthropus*, to delay fracture-related tooth loss (Kay, 1981; Lucas et al., 2008). Although chipping cannot conclusively be used to infer whether thick enamel in *Paranthropus* evolved to counter attrition or fracture, it seems chipping should not be used to support the hypothesis that thick enamel evolved to protect against fracture in *Paranthropus*. This is especially the case since molars are often the focus of such research, and in *P. robustus* a low chipping prevalence is evident in molars.

Other enamel characteristics also need to be considered with microstructure (e.g., Hunter-Schreger band thickness and enamel prism density) and overall tooth morphology potentially contributing to reduce fracture or limit chip size (Constantino et al., 2009). Further comparative studies can elucidate the functional and evolutionary implications of enamel structure in *Paranthropus*. The present study suggests that dental chipping in *Paranthropus* was rare relative to other hominins and extant durophagous primates. The ability of such enamel chips to deduce the selective pressures influencing the Paranthropines’ unique craniofacial hypodigm remains opaque. Additionally, differences in tooth properties among species, and how these differences influence fracture likelihood, need to be more adequately explored. Future work is needed to determine the causal factors for the low chipping frequency in paranthropines. However, based on extant primate comparisons, these chips should not be used as evidence to suggest *Paranthropus* regularly masticated hard foods.

## Acknowledgements

Research was supported by a Sir Thomas Kay Sidey Postdoctoral fellowship from the University of Otago to IT. We acknowledge Luke D. Fannin for helpful comments and suggestions on the manuscript. We also give thanks to the reviewers, associate editor, and co-editor-in-chief, whose comments greatly improved the manuscript.

